# Investigating postharvest seed coat darkening in common beans: a molecular perspective beyond the major *P* gene

**DOI:** 10.1101/2025.03.19.643989

**Authors:** Nishat Islam, Sangeeta Dhaubhadel

## Abstract

Many market classes of common beans (*Phaseolus vulgaris*) have a significant reduction in crop value due to the postharvest darkening of the seed coat. Seed coat darkening is caused by an elevated accumulation and oxidation of proanthocyanidins (PAs). In common bean, the major color gene *P* encodes for a bHLH protein with its *P^sd^*allele controlling the postharvest slow darkening seed coat trait. In the present investigation, we determined that P, PvMYB3A, and PvWD9 are the essential components that form a protein complex and regulate PA biosynthesis in pinto beans. P does not bind with the PA biosynthetic gene *Anthocyanin Reductase* (*PvANR*) promoter but regulates its expression by interacting with PvMYB3A, which directly binds to the *PvANR* promoter. PvWD9 is an essential member of the core protein complex, which requires one or more additional plant components in order to interact with its partner proteins P and PvMYB3A and create a functioning complex. The P^sd^ isoform affects the accumulation of PA by functioning in a similar manner to its isoform P, albeit at a lower efficiency. Understanding the regulation of PA biosynthesis in common beans helps to explain variances in seed coat color and issues associated with darkening after harvest.

**Highlight:** Color and pattern of the common bean seed coat are important traits in bean breeding, which is determined by level of proanthocyanidins (PA). This study provides experimental evidence for the regulation of PA biosynthesis in common beans by the P-PvMYB3A-PvWD9 complex together with some yet unknown associated protein(s). In the slow darkening beans, substitution of P with its slow dakening isoform P^sd^ reduces the target biosynthetic gene expression inflencing PA production.

## Introduction

Several major market classes of common beans (*Phaseolus vulgaris*), including pinto, carioca, cranberry, and pink, are affected by postharvest seed coat darkening. Bean producers and vendors face significant crop value losses due to decreased consumer preference for darker beans. Darker beans also exhibit poor canning quality and require longer cooking times (Park and Maga, 1999, Pirhayati *et al*., 2011, Cichy *et al*., 2019). While special storage conditions can somewhat slow down bean darkening, maintaining such conditions is expensive. There are at least three different postharvest darkening phenotypes in common bean: regular darkening (RD), slow darkening (SD), and non-darkening (ND), which are directly associated with the proanthocyanidins (PAs) levels in the seed coat (Elsadr *et al*., 2015, Freixas *et al*., 2017, Duwadi *et al*., 2018).

PA biosynthesis is generally determined by the expression of the late biosynthetic genes (LBGs) of the phenylpropanoid pathway, which include *Dihydroflavonol 4-Reductase (DFR)*, *Anthocyanidin Synthase (ANS)*, *Leucoanthocyanidin Reductase (LAR)*, and *Anthocyanidin Reductase (ANR)*. The expression of these genes begins just before seed development, specifically in flowers, and their expression peaks in young siliques (Nesi *et al*., 2001, Lepiniec *et al*., 2006, Li *et al*., 2016). Common bean seed coats with varying postharvest seed coat darkening traits exhibit differences in the accumulation of LBG transcripts (Figure 1).

**Figure 1.**
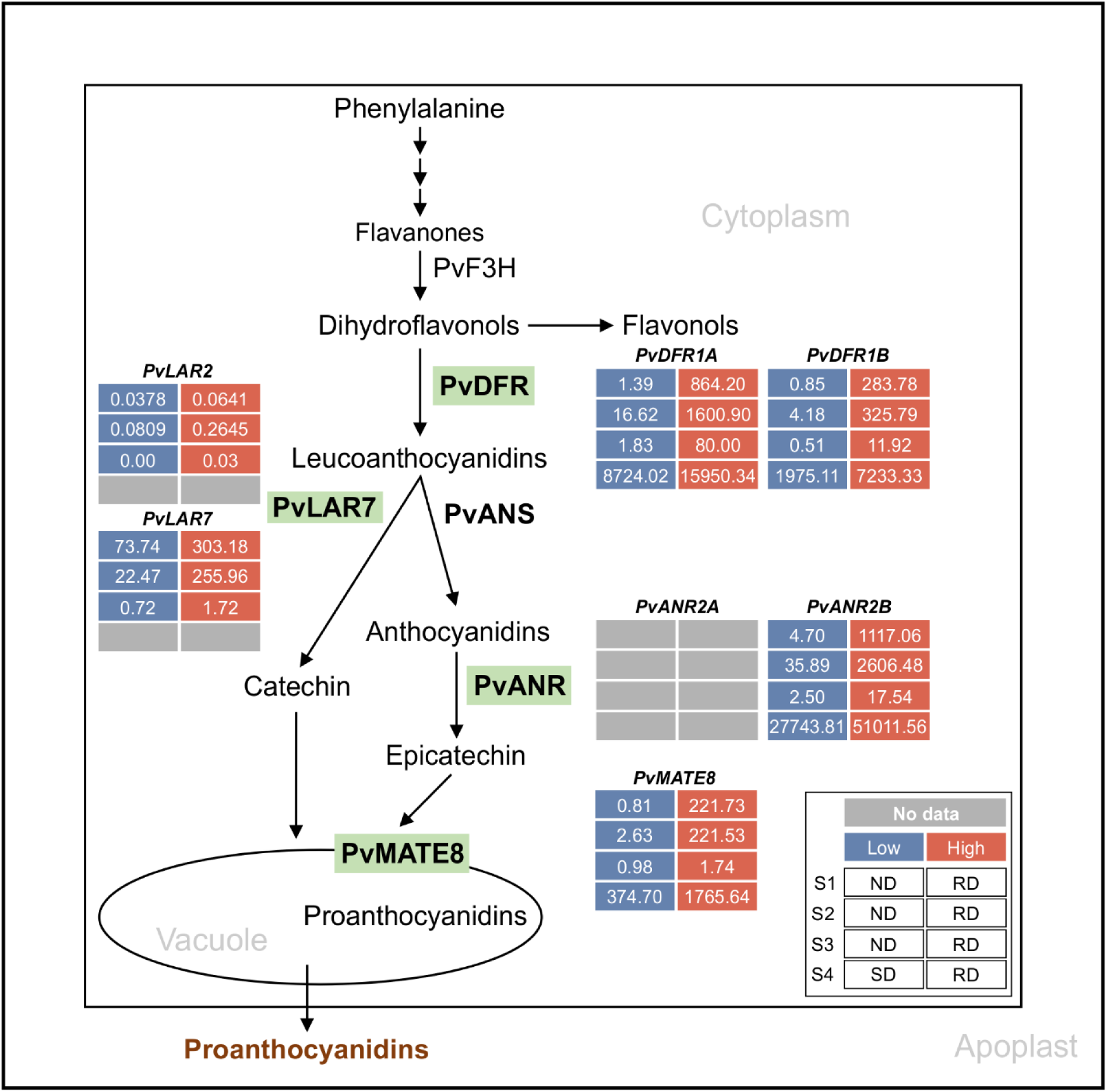
Proanthocyanidin-specific branch of phenylpropanoid biosynthetic pathway in common bean. Green highlights indicate the enzymes encoded by the late biosynthetic and transporter genes. Differential transcript expression in RD, SD and ND lines for the candidate genes are shown (legend on the inset). Inset description: S1, S2 and S3 indicate early, intermediate and mature stages of seeds and S4 represent the 50 mg seed coat used in Coutin et al., 2019 and Duwadi et al., 2018, respectively. Low, high and no available transcript values are color coded.

Complementation of pinto bean *P* in Arabidopsis *tt8* mutant activated *DFR* and *ANR* expression, and restored the wild type phenotype while its *P^sd^* allele only partially rescued the mutant phenotype (Islam *et al*., 2020). Pintos with P and P^sd^ display RD and SD seed coat phenotypes, respectively (Islam *et al*., 2020), while non-functional P produces a white color seed phenotype (McClean *et al*., 2018). The P protein belongs to the plant bHLH subgroup IIIf (Heim *et al*., 2003, Zimmermann *et al*., 2004) and contains a highly conserved 10-15 bp basic region of the bHLH domain responsible for DNA binding to the canonical E-box (CANNTG) or G-box (CACGTG) motif of the target gene promoters (Toledo-Ortiz *et al*., 2003, Feller *et al*., 2006, Hichri *et al*., 2011, Kong *et al*., 2012). *P* produces two transcript variants, where *P-1* is the major transcript responsible for PA accumulation in pinto beans (Islam *et al*., 2020). The protein encoded by *P-2* lacks the ACT domain at the C terminal end and it is not involved in PA biosynthesis.

In many plant species, the LBGs of PA pathway are regulated by the MYB-bHLH-WD40 (MBW) protein complex (Ramsay and Glover, 2005, Xu *et al*., 2015, Zhang and Schrader, 2017). In Arabidopsis, MYB (TT2), bHLH (TT8) and WD40 (TTG1) determine PA accumulation by regulating the expressions of *DFR* and *ANR* (Devic *et al*., 1999, Nesi *et al*., 2000, Nesi *et al*., 2001, Debeaujon *et al*., 2003). A similar mechanism also exists in *Medicago truncatula* (Dixon *et al*., 2005, Li *et al*., 2016), *Lotus japonicus* (Yoshida *et al*., 2008, Yoshida *et al*., 2010a, Yoshida *et al*., 2010b) and *Zea mays* (Grotewold *et al*., 2000, Hernandez *et al*., 2007). However, species-specific unique features are associated with the composition and function of the MBW complexes, which include the involvement of multiple MYB and bHLH members, protein-protein interaction patterns and priorities, and variation in their gene targets. For example, AtMYB5 (Gonzalez *et al*., 2009) and MtMYB14 (Liu *et al*., 2014) perform a similar function as AtTT2 and MtPAR, respectively. In persimmon fruit (*Diospyros kaki*), the WD40 (DkWDR1) interacts with the MYB (DkMYB2 and DkMYB4) partners but not with the bHLH (DkMYC1) (Gil-Muñoz *et al*., 2020). On the other hand, apple (*Malus domestica*) WD40 (MdTTG1) interacts with the bHLH (MdbHLH) but not with the MYBs (MdMYB9 or MdMYB11) in the MBW complex (An *et al*., 2014). Many species contain multiple homologs of MYB and bHLH proteins of the MBW complex, while only a single WD40 has been reported for all the plant species studied so far (Grotewold *et al*., 1994, Yoshida *et al*., 2008, Hichri *et al*., 2011, Islam and Dhaubhadel, 2023). Recently, possible MYB and WD40 partners of P and formation of protein complex have been suggested (McClean *et al*., 2024, Parker *et al*., 2024).

Here, we identified the protein partners of the P using an independent approach and validated the molecular mechanism of their action in common bean. We demonstrate that P is not a DNA binding protein. It interacts with two PvMYBs independently, and activates LBG expressions thereby influencing PA biosynthesis, albeit at different PA accumulation levels. Furthermore, interaction of P and/or PvMYB with PvWD40 requires additional plant protein(s). Formation of this proien complex is critical for the activation of LBGs and PA accumulation in common bean. Therefore, these findings will advance our understanding of the seed coat darkening mechanism and will assist bean breeders to address seed coat color-related concerns.

## Materials and Methods

### Plant material and growth conditions

Pinto bean cultivars CDC Pintium and 1533-15 were obtained from Dr. Kirstin Bett, University of Saskatchewan, Canada. *A. thaliana* ecotype Wassilewskija (WS2) and *tt2* mutant seeds (line: DROTV55T3) were obtained from the Institut Jean-Pierre Bourgin, Versailles, France, and *A. thaliana* ecotype Columbia-0 and *ttg1* mutant seeds were obtained from Arabidopsis Biological Resource Center (ABRC), Ohio State University, USA.

Surface sterilized pinto bean seeds were grown in Pro-Mix PGX soil under a light intensity of 300-400 μmol photons/m^2^/s. *A. thaliana* seeds were surface sterilized and grown on Murashige and Skoog medium for 1 week then transferred to the soil mix and grown at a light intensity of 150 μmol photons/m^2^/s. *Nicotiana benthamiana* plants were grown in Pro-Mix PGX soil under a light intensity of 80 μmol photons/m^2^/s. All plants were grown in a growth room in a 16 h light at 25°C and 8 h dark at 20°C cycle.

### In silico analysis

All genomic and proteomic data were retrieved from Phytozome v13 for *P. vulgaris* landrace, Chacha Chuga (G19833) (https://phytozome-next.jgi.doe.gov/info/Pvulgaris_v2_1). Multiple sequence alignments were carried out using Clustal Omega followed by BOXSHADE 3.21 (https://www.ch.embnet.org/software/BOX_form.html) (Sievers *et al*., 2011). Phylogenetic trees were built using neighbour-joining program with the bootstrap set to 1000 in MEGAX (Kumar *et al*., 2018). The phylogenetic tree was visualized by the iTOL online server (Letunic and Bork, 2021).

For promoter sequence analysis, a 2 kb upstream from the start codon (ATG) of *PvUANR2B* was obtained for *P. vulgaris* pinto UI111 1.1 genome from Phytozome v13 (https://phytozome-next.jgi.doe.gov/info/PvulgarisUI111_v1_1). Both strands of the DNA were used to search for potential MYB and bHLH binding sites prediction in New PLACE (https://www.dna.affrc.go.jp/PLACE/?action=newplace).

### RNA extraction and quantitative reverse transcription-PCR

Total RNA from bean seed coat was extracted using the RNeasy Plant Mini kit (Qiagen, USA). On-column DNA digestion was performed using DNase I (Promega, USA). cDNA was synthesized from total RNA (1.0 μg) using the SuperScript® IV Reverse Transcriptase (RT) (Invitrogen, USA). Platinum® Taq DNA Polymerase High Fidelity (Invitrogen, USA) was used for PCR. Quantitative real-time polymerase chain reaction (qRT-PCR) was performed using gene-specific primers (Supplementary Table 1) in three technical replicates for each sample using the SsoFast EvaGreen supermix (Bio-Rad) and CFX96 real-time PCR system (Bio-Rad). Gene expression was normalized to *PvUBQ* (*Phvul.007G052600.1*). Data analysis was conducted using CFX Maestro (Bio-Rad).

### Plasmid construction

For protein-protein interaction and gene overexpression, plasmids were constructed using Gateway technology (Invitrogen, USA). Genes of interest were cloned into Gateway destination vectors, pEarleyGate201-YN and pEarleyGate202-YC for BiFC (Lu *et al*., 2010), pEarleyGate101 for subcellular localization (Earley *et al*., 2006), pMDC32 for overexpression in Arabidopsis (Curtis, 2003), pK7WG2D for overexpression in hairy roots (Karimi *et al*., 2002) and pGADT7-GW for Y1H and Y2H as prey and pGBKT7-GW for Y2H bait (Clontech, USA) using Gateway primers (Supplementary Table 1). Overexpression of P-PvMYB3A and P-PvMYB10B gene fusions were created using an 18 bp linker (5’-AGCACAACATTTCAACCA-3’) by fusion PCR using the primers listed in Supplementary Table 1. For promoter analysis, DNA fragments were PCR amplified and sub-cloned into pGEM®-T easy vector by TA cloning (Promega, USA). Sequence confirmed promoter fragments were cloned using the *Hin*dIII and *Nco*I restriction sites of pGreenII-0800-LUC (Creative Biogene, USA) for dual-luciferase assay.

For plant transformations, plasmids were transformed into *Agrobacterium tumefaciens* strain GV3101 and for hairy root transformation, plasmids were transformed to *A. rhizogenes* strain K599 by electroporation. For Y1H and Y2H assays, recombinant plasmids were transformed to *Saccharomyces cerevisiae* strain Y1HGold (Clontech, USA) and AH109 (Clontech, USA), respectively using Frozen-EZ Yeast Transformation II Kit (Zymo Research, USA).

### Generation of common bean hairy roots

To generate hairy root transformants, 3-4 days old pinto bean (cultivar CDC Pintium) plants with unfolded green cotyledon were used following the method described by Estrada-Navarrete *et al*. (2007). Briefly, a single colony of *A. rhizogenes* strain K599 harboring genes cloned in the pK7WG2D vector was streaked on spectinomycin (100 μg/mL) containing LB plate. The bacterial inoculum was prepared by resuspending an overnight grown culture in 1 mL Milli Q water and 100 µM acetosyringone and incubated at room temperature for at least 1 hour with gentle agitation to activate the virulence genes required for transformation. Young plants were wounded just below the cotyledon and infected with 5 µL of the inoculum. Plants were covered with a plastic dome with increased humidity. In about 15 to 18 days, transgenic roots were screened for GFP expression, flash frozen and stored at −80°C until use.

### Subcellular Localization and Bimolecular Fluroecent Complementation (BiFC) assay

For subcellular localization, *A. tumefaciens* harboring gene of interest in pEarleyGate101 and nuclear marker were co-infiltrated in 1:1 (v/v) ratio into *N. benthamiana* leaves (Sparkes *et al*., 2006). For BiFC assay, pEarleyGate201-geneA-YN and pEarleyGate202-geneB-YC were co-infiltrated in a 1:1 (v/v) ratio into *N. benthamiana* leaves and visualized 48 h post-infiltration, using an Olympus FV1000 confocal microscope (Olympus Corporation, Japan) with a 60× water immersion objective lens. For YFP visualization, the excitation wavelength was set to 514 nm, and emission was collected at 530-560 nm. For CFP visualization, the excitation wavelength was set to 434 nm and emission was collected at 470-500 nm. To visualize the co-localization of the YFP and CFP signals, the ‘Sequential Scan’ tool was used.

### Genetic complementation of Arabidopsis

Arabidopsis plants were transformed with *A. tumefaciens* harbouring contructs with gene of interest using the floral dip infiltration method (Clough and Bent, 1998). T1 seeds were screened on MS media containing hygromycin (50 mg/L), and transformed plants were grown in soil to collect seeds for further analysis.

### Metabolite extraction, PAs quantification and staining

For quantitative analysis of PAs, 0.02-0.03 mg of hairy roots were extracted with 10 volumes of 50% acetonitrile (Sigma-Aldrich, Canada). Samples were sonicated for 20 min at 4°C followed by centrifugation at 11,200× *g* for 10 min at 4°C. Supernatants were collected and used for spectrophotometric assay as described by Coutin *et al*. (2017). Absorbance was measured at 640 nm using Synergy^TM^ 2 Multi-Detection microplate reader (BioTek). Proanthocyanidin A2 was used as a standard (BOC Sciences, USA).

Arabidopsis seeds were stained with 0.5% (w/v) 4-dimethylamino-cinnamaldehyde (DMACA) (Sigma-Aldrich, Canada) for 36 h in dark followed by de-staining with 70% ethanol (Li *et al*., 1996, Zhao and Dixon, 2009). For seed mucilage observation, 50 mM EDTA was added to approximately 20 mature and desiccated Arabidopsis seeds, and incubated at room temperature in a shaker at 15× *g.* After 2 h, the solvent was removed completely from the seeds, and 0.01% ruthenium red (Sigma-Aldrich, Canada) solution was added and placed on a shaker at 15× *g* for another hour. Seeds were washed with Milli Q water and photographs were taken using an SMZ1500 dissecting microscope with camera (Nikon).

### Transient expression and dual-luciferase assay

*A. tumefaciens* GV3101 culture containing reporter and effector plasmids were mixed in a 1:9 ratio and infiltrated into *N. benthamiana* leaves. Firefly Luciferase (LUC) and *Renilla* Luciferase (REN) activities were measured using the dual-luciferase assay reagents kit following manufacturer’s instructions (Promega, USA). Absolute relative luminescence unit (RLU) was measured as a ratio of LUC/REN. Plant transformed with reporter without any effector was used as a control.

### Yeast one-hybrid (Y1H) and two-hybrid (Y2H) assays

For Y1H assay, a −1630 bp region of *PvUANR2B* cloned in pAbAi vector were linearized with *Bst*B1 before transforming into *S. cerevisiae* Y1HGold strain (Clontech, USA) to integrate the promoter as a bait fragment into the yeast genome. Thereafter, the transformants were cultivated in culture plates on SD/-Ura media at 30°C for 3 days. The transformed colonies were PCR screened using Matchmaker Insert Check PCR Mix1 (Clontech, USA). The positive colonies were used for competent cell preparation using Frozen-EZ Yeast Transformation II Kit (Zymo research, USA). Y1H assays were performed by following the Matchmaker Y1H user manual (Clontech, USA). The minimum inhibitory concentration of Aureobasidin A (AbA) for yeasts carrying the promoter baits was 100 ng/mL. The prey constructs in pGADT7 vector were transformed into yeast carrying a promoter bait fragment, plated on SD/-Leu/AbA^150^, and incubated at 30°C for 3-5 days to observe the interaction.

For Y2H assays, *S. cerevisiae* strain AH109 was transformed with a 1:1 mixture of bait (pGBKT7-geneA) and prey (pGADT7-geneB) plasmids using the Frozen-EZ Yeast Transformation II Kit (Zymo Research, USA) and selected on SD/-Leu/-Trp agar medium. Selected individual yeast transformants were grown in liquid medium and a 5 µL of the culture was spotted on onto SD/–Leu/–Trp and SD/–Ade/–His/–Leu/–Trp plates for 3 days at 30°C.

## Results

### P is necessary but not sufficient for PA biosynthesis in common bean

Previously we demonstrated that P (the P-1 isoform) is able to restore the regulatory function of TT8 in Arabidopsis *tt8* mutant line (Islam *et al*., 2020). To dissect the mechanism how P regulates PA biosynthesis in common bean, we aimed to check the expression of late biosynthetic genes (LBGs) involved in PA biosynthesis. For this, the LBGs needed to be identified first in common bean. Thus, BLASTp searches were conducted with the protein sequences of *M. truncatula* DFR (Accession Q6TQT1), MtANR (Accession Q84XT1) and MtLAR (Accession CAI56327) queried against the *P. vulgaris* proteome database in Phytozome v13. The search resulted in 26 PvDFR and 7 PvANR candidates, which were used to build phylogenetic trees with characterized DFR and ANR proteins from other plant species (Supplemental Figure 1). Based on the phylogenetic analysis, Phvul.001G012700 (PvDFR1A), Phvul.001G012800 (PvDFR1B), and Phvul.007G246200 (PvDFR7) were found to cluster with DFRs and Phvul.002G218600 (PvANR2A) and Phvul.002G218700 (PvANR2B) with ANRs from other plant species that have been characterized. The search returned Phvul.007G102100 (PvLAR1) with a 78% and Phvul.002G033300 (PvLAR2) with 42% identity to MtLAR at the amino acid level.

The tissue-specific expression of the shortlisted candidates were analyzed using publicly available RNAseq data for common bean and heatmaps were generated (Figure 2a). Since PA is a seed coat-specific trait in common bean, emphasis was given to the seed-related tissues for the expression of the target genes. *PvDFR1A,* and *PvANR2B* showed the highest expression in young pods (67.12 and 97.68, respectively) and *PvDFR1B* showed the highest expression in young pods (28.71) and flower buds (28.71) expressed in Fragments Per Kilobase of transcript per Million mapped reads (FPKM). Additionally, *PvDFR7* was mostly expressed in roots but only a negligible amount was present in seeds, therefore, it was not included in further analyses. In spite of its tissue-specific expression, *PvANR2A* was included in the analysis for the possibility of being a paralogue of *PvANR2B* (Supplemental Figure 1b). Among *PvLAR1* and *PvLAR2*, the later showed high expression in the root tissues and a negligible about for seed-related tissues, thus not included in the analysis.The *PvLAR1* expression was highest in the flower buds (98.52). The transporter *PvMATE8* showed the highest expression in young trifoliate (21.83), followed by flowers (10.74).

**Figure 2.**
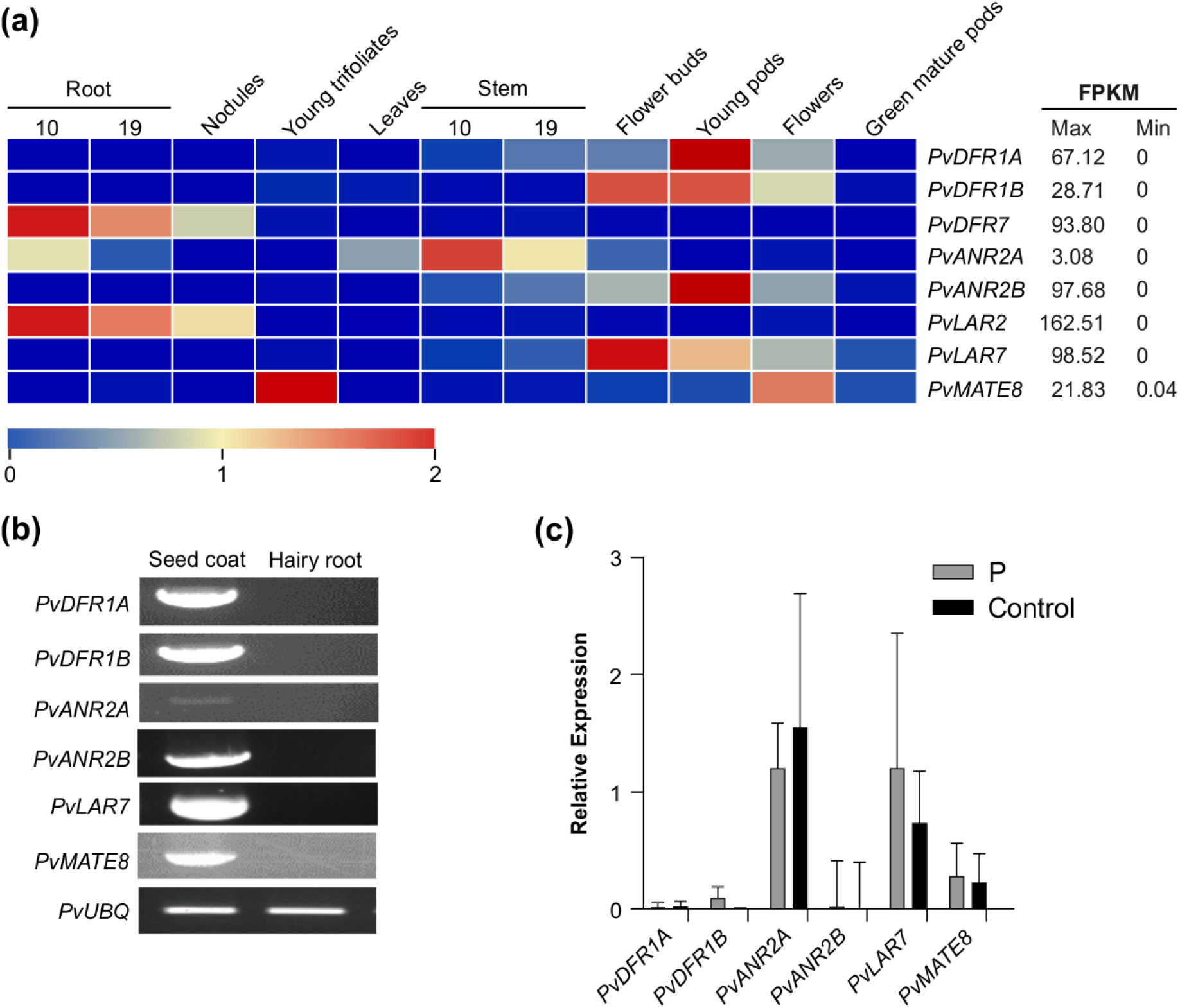
Expression of candidate PA biosynthetic and transporter genes in different tissues of common bean. (a) tissue-specific expression profile of the target genes in common bean. Transcript data retrieved from Phytozome for landrace G199833 was used for the generation of heatmap. The color key represents the relative transcript abundance in a row with the log_2_ transformed values. (b) RT-PCR showing expression of candidate genes in pinto bean seed coat and hairy root tissues. (c) RT-qPCR showing the transcript abundance of PA biosynthetic and transporter genes in the hairy root lines overexpressing *P* and control.

Since plant transformation and *in vitro* regeneration system in common bean is not efficient and reproducible (Mukeshimana *et al*., 2013), we used hairy root system as an alternative approach to validate gene function. Additionally, absence of the LBGs and PA transporter *PvMATE8* expression in root tissue (Figure 2b) provides additional advantage as hairy roots could serve as mutants in studying PA biosynthesis in beans. Pinto bean cv CDC Pintium hairy roots overexpressing *P* were generated and transgenic roots were screened using GFP expression. Control hairy roots were generated by using the *A. rhizogenes* K599 with an empty vector. Hairy roots were obtained in 15 to 18 days after transformation with around 60% transformation efficiency (Supplemental Figure 2). Transgenic roots overexpressing *P* gene were assessed for both the alteration in the expression of the LBG targets and the total PA levels. The results showed no differences on the transcript levels of *PvDFR1A*, *PvDFR1B*, *PvANR2B* and *PvMATE8* in *P* overexpressed and control hairy root lines (Figure 2c). *PvANR2A* and *PvLAR1* transcripts were present in the control roots and their levels were not altered significantly in the transgenic hairy roots overexpressing *P*. Furthermore, no PA was detected in the *P* overexpressing lines. Based on the transcript and metabolite analyses, we conclude that the P alone is not sufficient for PA biosynthesis in pinto bean.

### A search for P interactome identifies two MYBs and a WD40 protein involved in PA biosynthesis in pinto bean

In many plants, MBW complex regulates the expression of the structural genes such as *DFR*, *ANR*, *LAR,* and *MATE* transporter in the PA biosynthetic pathway (Xu *et al*., 2014). Since P alone was not adequate for PA LBG activation, we searched for its MYB and WD40 partners that potentially interact with P to complete the regulatory function of PA biosynthesis.

To identify the potential interactors of P, BLASTp searches using protein sequence of MtPAR (Accession ADU78729) and MtWD40-1 (Accession ABW08112.1) as queries were performed against the *P. vulgaris* proteome, followed by phylogenetic and transcript abundance analysis in seed forming tissues, and identified two candidate PvMYBs, Phvul.003G132100 (PvMYB3A) and Phvul.010G130500 (PvMYB10B) and two candidate PvWD40s, Phvul.001G098000 (PvWD1A) and Phvul.009G044700 (PvWD9) (Supplemental Figures 3 and 4). We also confirmed that the candidate PvMYBs and PvWD40s are localized in the nucleus (Supplemental Figure 5). To investigate whether P interacts with the candidate PvMYBs and PvWD40s, *in vivo* protein-protein interaction assays were performed by BiFC assay. The interaction was visualized using confocal microscopy where presence of fluorescence indicated direct or close interaction as well as subcellular location of interaction. Our BiFC results revealed that P interacts with both PvMYB3A and PvMYB10B, and with PvWD9 in the nucleus (Figure 3a). P and PvWD1A interaction was not detected, therefore, PvWD1A was excluded from further analysis. Furthermore, PvWD9 interacted with both PvMYB3A and PvMYB10B. No fluorescence was observed between the constructs containing genes of interests and empty vector controls containing the complementary YFP fragments. Similar results were obtained when BiFC assays were performed using reciprocal combinations (Supplemental Figure 6).

**Figure 3.**
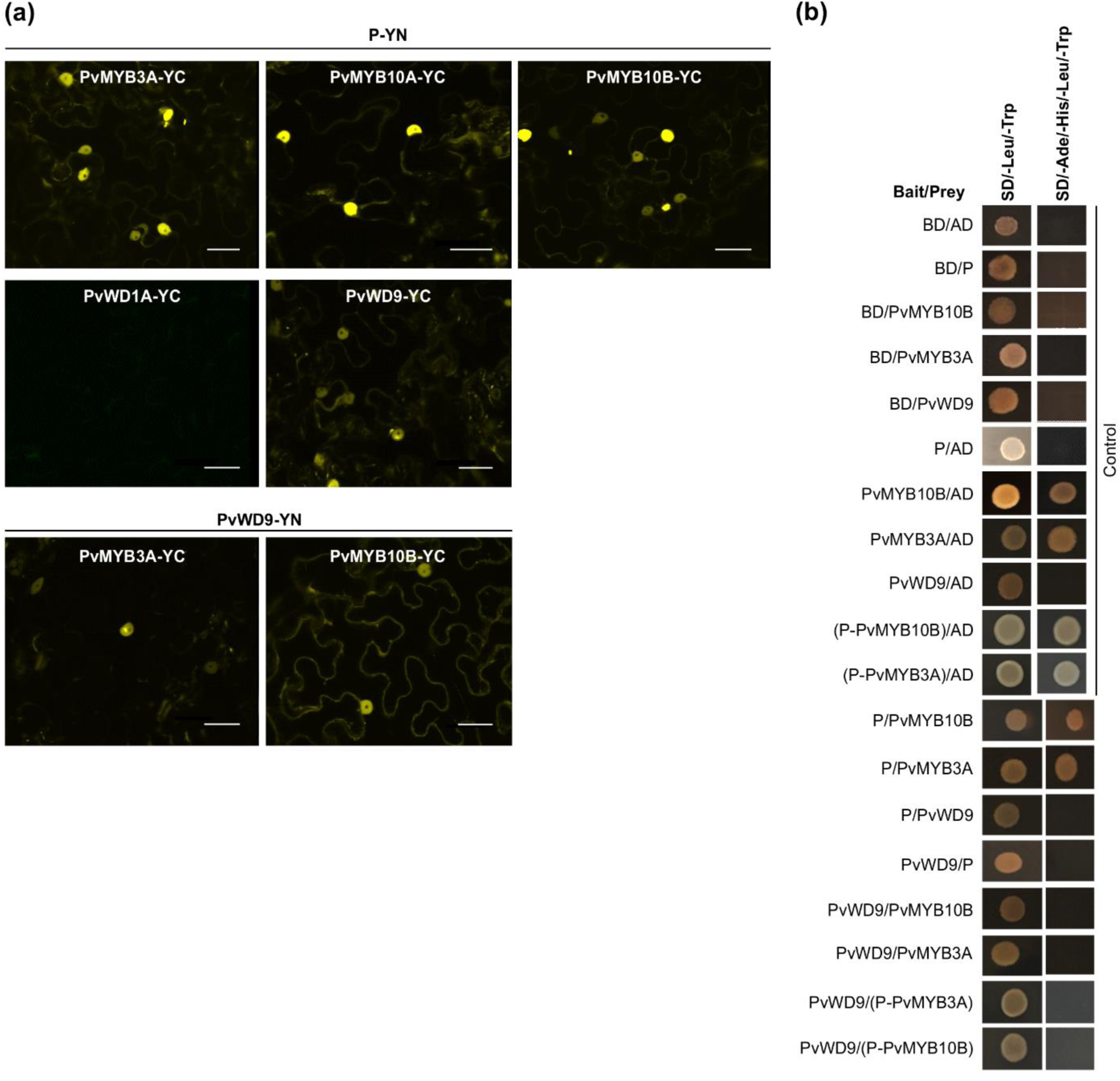
Protein-protein interaction between candidate proteins involved in MBW complex formation. (a) BiFC assay between proteins was assayed by co-expression of translational fusions with N–terminal (YN) and C–terminal (YC) fragments of YFP in *N. benthamiana*. Proximity of the two fragments results in a functioning fluorophore. Scale bar= 40 µM. (b) Yeast cells were co-transformed with combination of DNA-binding domain (BD, Bait) and activation domain (AD, Prey) fused constructs as indicated. Yeast suspension culture (5 μL) was spotted onto synthetic defined (SD) selection plates. Growth on SD plates without leucine and tryptophan (SD/-Leu/-Trp) showed the presence of both the vectors, while growth on SD plates without leucine, tryptophan, adenine and histidine (SD/-Ade/-His/-Leu/-Trp) indicated interaction between bait and prey.

**Figure 4.**
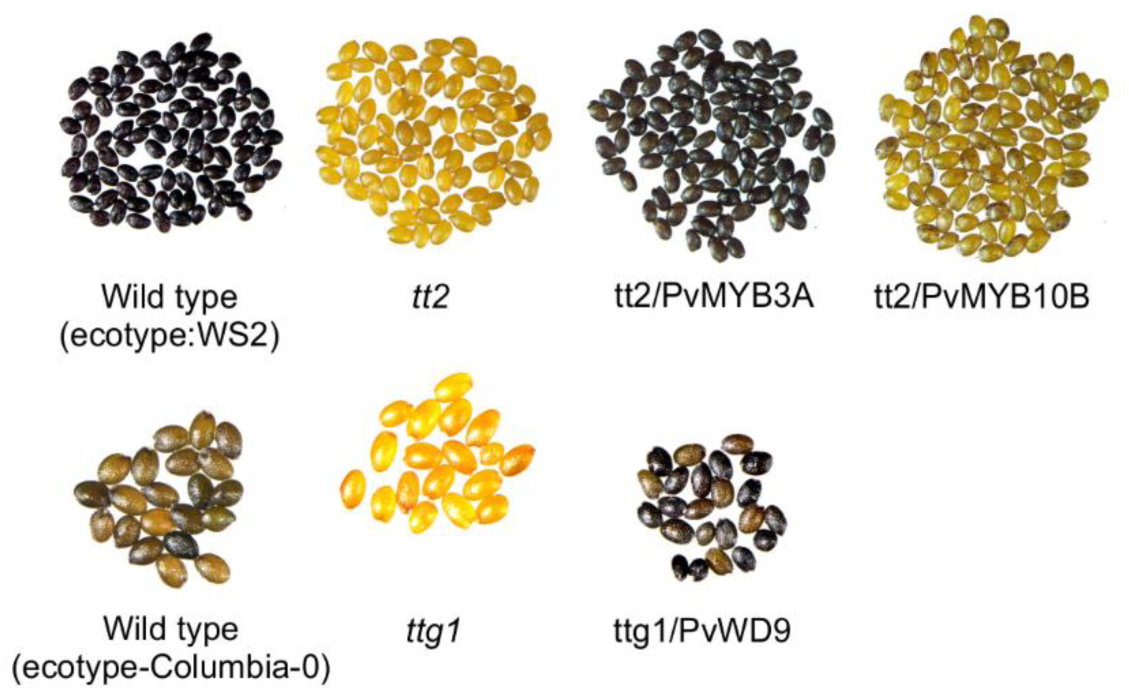
*PvMYB3A* and *PvWD9* rescued Arabidopsis *tt2* and *ttg1* mutant phenotypes. Seeds of *WS2, tt2* and *tt2* transformed lines with PvMYB10B and PvMYB3A (top). Seeds of columbia-0, *ttg1* and *ttg1* transformed with PvWD9 (bottom). All seeds were stained in DMACA reagent for 36 h, washed in 70% ethanol and picture taken under light microscope.

To futher confirm the protein-protein interaction, interaction assays between P, PvMYBs and PvWD9 in pairwise combinations were performed by using a targeted Y2H assay. All the candidate proteins were independently used as either a bait protein (fused with DNA-binding domain, BD) or a prey protein (fused with activation domain, AD) (Figure 3b). Controls included each of the test proteins as bait with an empty vector as prey and its reciprocal combinations. Additionally, a combination of both empty vectors (BD/AD) was also included as a negative control. Controls were included in each experiment on the same plate to ensure the plate quality.

As shown in Figure 3b, both PvMYB3A and PvMYB10B (baits) and empty vector (prey) co-transformed yeast colonies grew on the synthetic defined (SD) media lacking Ade, His, Leu and Trp suggesting the auto-activation of PvMYB3A and PvMYB10B. Therefore, Y2H assays were performed in all combinations of baits and prey except for the ones that showed auto-activation. The results revealed that P interacts with PvMYB3A and PvMYB10B but not with PvWD9. Furthermore, PvWD9 did not interact with either of the PvMYBs in the Y2H assay. Since PvWD9 did not interact with P and both PvMYBs in yeast, we speculated that both P and MYB together are possibly required for PvWD9 binding in the ternary complex. Thus, two separate fusion constructs were created by fusing *P* with *PvMYB3A* and *P* with *PvMYB10B* using an 18 bp linker. Both the constructs were cloned to the bait and prey vectors, and Y2H assays were performed with PvWD9. P-PvMYB3A and P-PvMYB10B when fused to the BD, showed auto-activation. In the Y2H assays for the fusion proteins, no interaction was observed between P-PvMYB3A and P-PvMYB10B with PvWD9 (Figure 3b). These findings suggest that some plant-specific factor(s) are possibly involved in the complex formation that allowed PvWD9 to interact with PvMYB3A, PvMYB10B and P in *N. benthamiana*, and that the factor was not present in yeast.

### PvMYB3A and PvWD9 restore PA biosynthesis in Arabidopsis mutants

Arabidopsis *tt* mutant lines serve as an efficient molecular tool for PA pathway gene characterization (Appelhagen *et al*., 2014). When TT2 (MYB) and TTG1(WD40) in the Arabidopsis MBW complex are non functional, PA biosynthesis in the seed coat gets abolished giving the *tt2* and *ttg1* mutant seeds pale yellow appearance (Nesi *et al*., 2001). To determine if PvMYB3A, PvMYB10B and PvWD9 regulate PA biosynthesis, we overexpressed the genes in their mutant background. T3 seeds were collected from multiple independent transgenic lines and phenotyped by DMACA staining. A representative photograph of stained seeds are shown in Figure 4 demonstrating that PvMYB3A and PvWD9 restored the wildtype phenotype in the complementation line. tt2/PvMYB10B seeds showed PAs as spots on the yellow background. Arabidopsis *TTG1* is expressed ubiquitously in all tissues and regulates five cellular functions: trichome formation, seed mucilage formation, root hair patterning, anthocyanin and PA accumulation by interacting with different MYB and bHLH proteins in Arabidopsis (Zhang and Schrader, 2017). PvWD9 restored all five TTG1 dependent traits in T2 lines (Supplemental Figure 7). These results confirm that PvMYB3A and PvWD9 play role in PA biosynthesis in Arabidopsis.

### P interacts with PvMYB3A to regulate PA biosynthesis

To determine the function and molecular mechanism of P, PvMYB3A, PvMYB10B and PvWD9 in common bean, we checked the expression of these genes in hairy roots. As shown in Figure 5a, *PvWD9* expression was detected in both seed coat and hairy roots, whereas *P*, *PvMYB3A* and *PvMYB10B* transcripts were present only in seed coat. Since PvWD9 transcripts were present in the roots, we speculated that, if the correct combination of the transcription factors are introduced in the hairy roots they would activate biosynthetic gene expression and subsequently the PA accumulation. Thus, *P*, *PvMYB3A* and *PvMYB10B* genes were introduced individually and as fusions of *P* and *PvMYB3A*, and *P* and *PvMYB10B* to generate transgenic hairy roots in pinto bean. The transgenic hairy roots were selected with GFP fluorescence and further verified for transgene expression by RT-PCR (Figure 5b). Once the transgenic nature of the hairy roots were confirmed, we performed quantitative analysis of LBG transcripts and PA levels. As shown in Figure 5c, *PvANR2A* transcript levels in the transgenic roots were not affected, indicating that *PvANR2A* is not the target of the MBW proteins used in this study. On the other hand, *PvANR2B* transcript levels were significantly increased in the transgenic roots overexpressing *P* (5.8 fold) and *P-PvMYB10B* (45.85 fold). A highly significant increase in the expression of *PvANR2B* was observed in P-PvMYB3A roots (456.98 fold) compared to the control. Similarly, a high accumulation of *PvDFR1A* (65.45 fold), *PvDFR1B* (560 fold) and *PvMATE* (8.5 fold) transcripts were found in P-PvMYB3A transgenic roots compared to control. A significant increase in the accumulation of *PvDFR1B* (95.90 fold) transcripts was also observed in P-PvMYB10B roots. Unlike the expression of other biosynthetic gene candidates, *PvLAR1* transcripts were high in *PvMYB3A* overexpressing (3.1 fold) roots suggesting that PvMYB3A alone is sufficient for *PvLAR1* gene activation.

**Figure 5.**
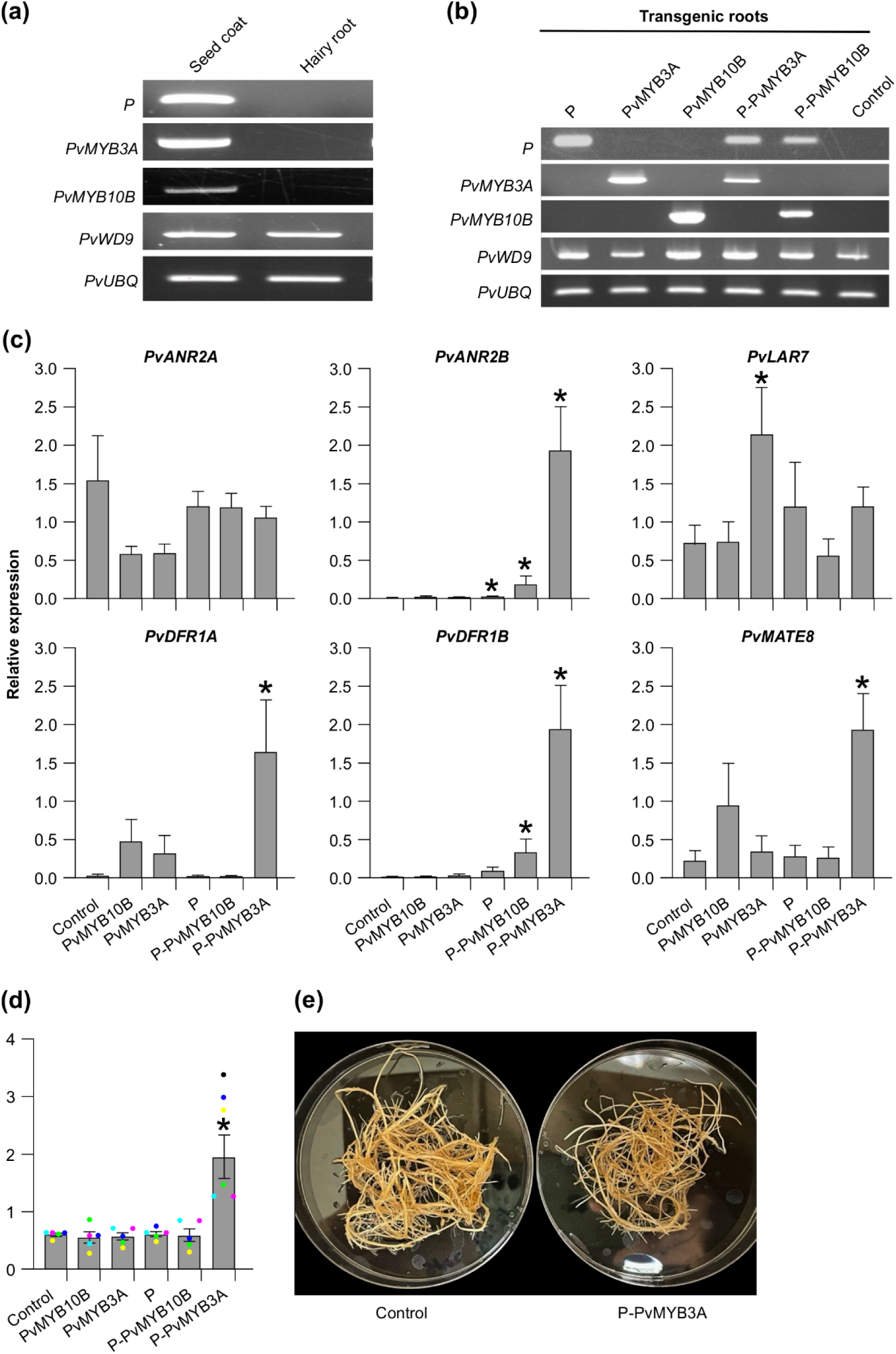
Transcript abundance and PA levels in the transgenic hairy roots of pinto bean. (a) and (b) RT-PCR showing expression of candidate MBW complex genes in seed coat and hairy root tissues of pinto bean cv CDC Pintium and confirmation of the transformation of respective hairy root lines with each of the constructs. *P. vulgaris* ubiquitin (*PvUBQ*) was used as reference gene control. (c) RT-qPCR was performed using gene-specific primers to check the expression of biosynthetic and transport gene candidates on each of the transgenic hairy root lines. Values were normalized with the reference gene *PvUBQ.* Empty vector transformed hairy root lines were used as control. Error bars indicate ±SEM (n=4, independent biological replicates and three technical replicates for each biological replicate). Asterisks (*) represent a significant difference compared to vector-only control as determined by Student’s *t*-test (p < 0.05). (d) Bars represent the total extractable PA levels expressed as proanthocyanidin A2 equivalents. Data represent the mean *±* SE (n=5, independent biological replicates and three technical replicates for each biological replicate). Asterisks (*) represent a significant difference compared to control as determined by Student’s *t*-test (p < 0.05). (e) Images of P-PvMYB3A overexpressing transgenic and control hairy roots.

Since the transgenic hairy root lines showed alteration in the transcript levels of PA biosynthetic and transporter genes, we evaluated how it affected PA levels. Among all transgenic roots, only P-PvMYB3A roots accumulated significantly higher levels of PAs compared to control (Figure 5d). Despite the difference in the PA levels, P-PvMYB3A hairy root line looked similar to control roots under white light (Figure 5e). Even though significant increases in the transcript levels of *PvANR2B* and *PvDFR1B* in *P-PvMYB10B* overexpressed roots were observed, PA levels remained unaltered and was same as the control (Figure 5c and 5d). Based on these results, we conclude that P requires PvMYB3A for the regulation of PA biosynthesis in common bean.

### PvMYB3A interacts with the promoter of *PvANR2B* and activates gene expression

To illustrate how P, PvMYB3A and PvMYB10B regulate PA biosynthetic gene expression, a promoter analysis was performed using *PvANR2B* gene promoter. Even though the coding gene sequences of *PvANR2B* in *P. vulgaris* G19833 and pinto UI111 cultivars are identical, major sequence variation in their promoter regions was observed (Supplemental Figure 8). Therefore, 2 kb upstream of the translation start site of *PvUANR2B* from *P. vulgaris* cultivar pinto UI111 were searched for *cis*-regulatory motifs. A total of 23 MYB and 4 bHLH binding motifs in the *PvUANR2B* promoter were identified (Supplemental Table 2).

To assess the DNA binding ability of P, PvMYB3A and PvMYB10B, a Y1H assay was performed using the 1630 bp upstream region from the translation start site of *PvUANR2B* (*PvUANR2Bpro*) from pinto bean cv CDC Pintium as the bait (Figure 6a). The result revealed that only PvMYB3A can directly bind with the *PvUANR2Bpro*. Despite the activation of *PvUANR2B* gene expression (Figure 5c) and presence of DNA-binding domains, neither P nor PvMYB10B showed DNA-protein interaction with the *PvUANR2Bpro*. Previously reported soybean GmCHS8-30bpTR and GmMYB176 (Anguraj Vadivel *et al*., 2021), and empty prey vector (pGADT7) were used as the positive and as negative controls, respectively.

**Figure 6.**
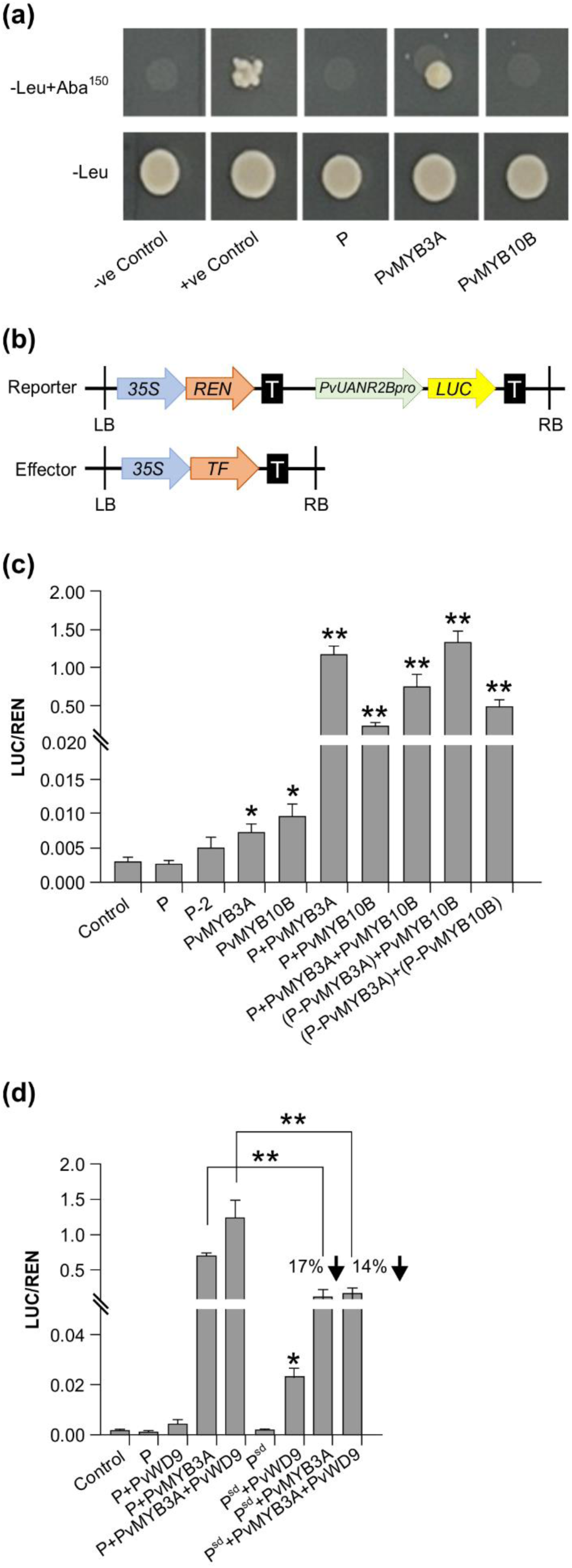
Effect of interactions between MBW proteins in different combinations on *PvUANR2B* promoter. (a) The Y1H Gold strains successfully transformed with the bait and prey vectors grown on SD/-Leu/AbA plates for 3-5 days at 30 °C. The interactions were confirmed using the yeast cell growth status. Positive control soybean GmCHS8-30bpTR and GmMYB176 (Anguraj Vadivel 2021) and negative control, pGADT7 empty vector were spotted on the same plate. (b) Schematic representation of reporter and effector constructs used in promoter transactivation assay. (c) and (d) Transient expression assays were performed by co-transforming reporter and effector plasmids at a molar ratio of 1:9 into *N. benthamiana* leaf. Relative luminescence was measured as a ratio of LUC/REN for each effector and their combinations. Reporter without any effector was used as a control. Data are presented as mean ± SE (n = 6, independent biological replicates and three technical replicates for each biological replicate). Asterisks denote significant difference relative to reporter-only (no effector) control as determined by Student’s *t*-test (* p<0.05, ** p<0.001).

The *PvUANR2Bpro* (−1630 bp) was cloned in a dual-luciferase vector to drive Firefly *Luciferase* (*LUC*) reporter gene expression (Figure 6b). The vector also contained a *Renilla Luciferase* (*REN*) reporter under the control of a CaMV 35S promoter in a separate cassette which was used to normalize the data. For the DNA-protein interaction assays, P, P-2 (P without the ACT domain), PvMYB3A, PvMYB10B, and P-PvMYB3A and P-PvMYB10B fusions were used as effectors. Both the reporter and effector clones were co-infiltrated in *N. benthamiana* leaves for transient expression and the binding of effector to the promotor region was monitored by measuring the luminescence. The relative expression was measured by determining the LUC to

REN ratio for each effectors (P, P-2, PvMYB3A or PvMYB10B) individually or their combinations (P+PvMYB3A, P+ PvMYB10B or P+PvMYB3A+PvMYB10B) or combinations with the fusion proteins [(P-PvMYB3A)+PvMYB10B and (P-PvMYB3A)+(P-PvMYB10B)]. As shown in Figure 6c, PvMYB3A and PvMYB10B signicantly increased *LUC* expression in comparison to the reporter-only control (no effector). Combination of P and PvMYB3A produced the highest accumulation of *LUC* transcripts (384 fold), whereas P and PvMYB10B produced a 71 fold increase of the same (Figure 6c, 7a, 7b). While *LUC* expression was greatly boosted (244 fold, compared to control) by the three transcription factors P, PvMYB3A, and PvMYB10B together (Figure 6c, 7d), it was not as much as when P and PvMYB3A were used as effector combinations (Figure 6c, 7a). Interestingly, *LUC* expression was 430 times higher than control when P-PvMYB3A fusion construct and PvMYB10B were utilized as the effectors, much like when P and PvMYB3A were employed as the effectors (Figure 6c, 7a, 7c). These findings point towards PvMYB3A as the primary MYB partner of P that is necessary for the PA biosynthetic gene expression. In addition, when P-PvMYB10 and P-PvMYB3A fusion proteins were introduced together, it showed *LUC* expression similar to that of P, PvMYB3A and PvMYB10B combined effectors (261 fold, compared to control) (Figure 6c, 7d, 7e). This indicates P does not exhibit preferential interaction with PvMYB10B or PvMYB3A, and that its binding partner dictates PA biosynthetic gene expression.

**Figure 7.**
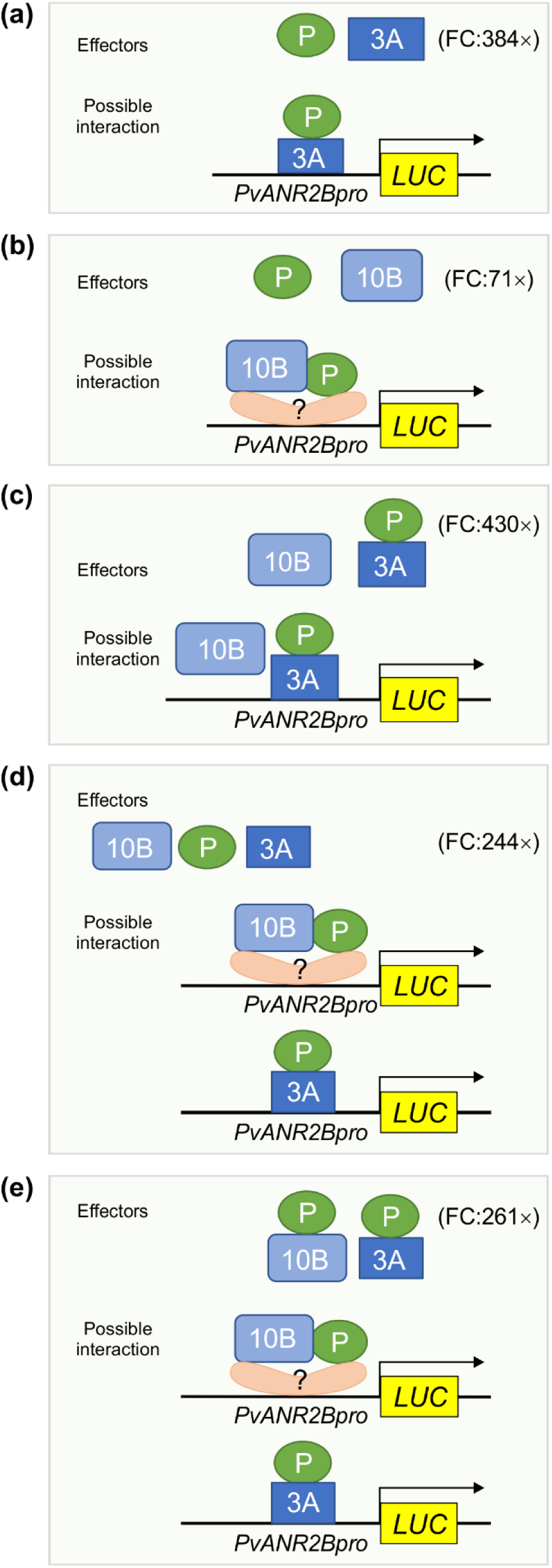
Proposed models showing the function of MBW complex partners for *PvUNAR2B* promoter driven gene activation. Based on the Y1H assay, only PvMYB3A (3A) but not P, and PvMYB10B (10B) binds with the *PvANR2Bpro*. Transient expression of *LUC* driven by the *PvANR2Bpro* demonstrated that P and 3A, either in combination as effectors (a) or as a fusion protein (c) can maximize *LUC* expression relative to the control, as shown by the fold change (FC). Despite the fact that 10B and P did not exhibit *PvANR2Bpro* binding ability, a 71-fold increase in *LUC* expression was seen (b) in comparison to the control, when they were utilized together as separate effectors. This suggests that 10B and P require the presence of co-factor(s) having DNA binding activity in order to stimulate gene expression. In the absence of P, 10B does not have significant effect on gene expression (c). Furthermore, P does not have any preferential interaction with 3A or 10B for *PvANR2Bpro* driven *LUC* expression (d, e).

**Figure 8.**
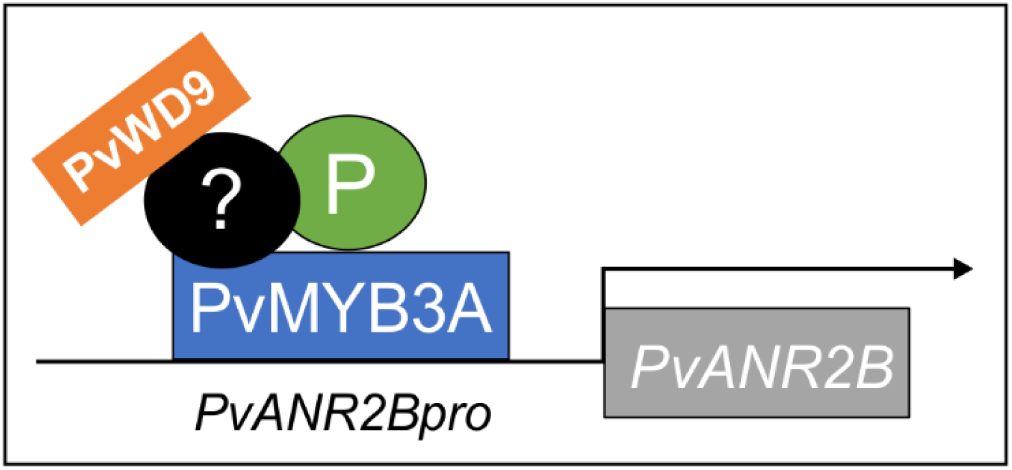
Infographic on *PvANR2B* gene activation by the MBW complex in pinto bean. P, PvMYB3A and PvWD9 along with additional factor(s) form a protein complex to activate late PA biosythetics genes (here, *PvANR2B*) thus regulating the synthesis and accumulation of PA in pinto bean seed coat. PvMYB3A only diectly interacts with the gene promoter.

To evaluate the role of PvWD9 in the protein complex and the impact of the protein complex in seed phenotype, LUC assay was performed using additional effectors-PvWD9 and P^sd^ (slow darkening isoform of P) with P, PvMYB3A in different combinations (Figure 6d). Our result revealed that P and PvWD9 together did not alter *LUC* expression relative to control. Whereas, PvWD9, P and PvMYB3A together increased *PvUANR2B* promoter activation significantly (636 fold) than in the P and PvMYB3A combination (361 fold) in *N. benthamiana* leaves, confirming that PvWD9 is a significant member of the protein complex. A 17% reduction in *LUC* expression with PvMYB3A and 14% reduction with PvMYB3A and PvWD9 together was found when P was replaced with its slow darkening isoform P^sd^ in the transactivation assay (Figure 6d). Thus, we conclude that P is not a DNA binding protein, but interacts with PvMYB3A and PvWD9 to form the complex where PvMYB3A binds directly with the promoter of *PvANR2B*.

## Discussion

PAs are well-known for their diverse functions in plants, which include protecting against predators and UV damage, boosting astringency and flavor of beverages, increasing the value of fodder crops, and conferring human health benefits (Dixon *et al*., 2005). In common beans, PAs contribute to the phenotypic variations across different market classes and also influence the postharvest appearance of seeds. We previously determined that P contributes to the postharvest seed coat darkening in pinto beans (Islam *et al*., 2020). Given that the orthologs of P function in a MBW complex in other plant species, and that different types of protein-protein interaction have been reported (Li *et al*., 2016, Lu *et al*., 2021, Xu *et al*., 2021), it is necessary to gain a deeper understanding of the molecular mechanisms behind seed coat darkening in pinto bean. Through the genome and transcriptome analysis, as well as interaction assays, here we identified two MYBs (PvMYB3A and PvMYB10B) and a WD40 (PvWD9) as potential interactors of P. Recently, two separate genetic investigations (McClean *et al*., 2024, Parker *et al*., 2024) reported *PvMYB3A* and *PvWD9* as *Z* and *T* genes, respectively, and *PvYB10B* as *J* gene (Erfatpour and Pauls, 2020). Our study demonstrates that P, PvMYB3A, PvMYB10B and PvWD9 interact with one another *in planta* (Figure 3a). The interaction of P with PvMYB3A and PvMYB10B was also confirmed by Y2H assay. However, the interaction of PvWD9 with P and PvMYB partners could not be validated in the yeast system (Figure 3b) indicating the involvement of additional plant factor(s) in the complex formation. It is more likely that the role of unidentified components is to provide a docking site necessary to form an active protein complex. In other plants, there are indications of the existence of additional interactor proteins. For example, in Arabidopsis, the WRKY transcription factor TTG2 interacts with the MBW complex to regulate seed-specific functions (PA biosynthesis and morphogenesis) and trichome formation (Johnson *et al*., 2002). Similarly, the WIP-type zinc finger TT1 (Appelhagen *et al*., 2011) and the MADS domain protein TT16 (Nesi *et al*., 2002) interact with the TT2 targeting particular steps of the flavonoid pathway.

PA biosynthesis occurs only in specific tissues such as seeds, fruits, and occasionally in the bark of trees but not in roots. Therefore, it is not anticipated that the genes involved in the PA biosynthesis will be expressed in roots (Nesi *et al*., 2000, Nesi *et al*., 2001, Dixon and Sarnala, 2020). The expression of regulatory (*PvMYB3A, PvMYB10B* and *P*) and PA biosynthetic (*PvDFR1A, PvDFR1B, PvANR2B, PvLAR1* and *PvMATE8*) genes is either very low or not present in root tissues, as shown in Figures 2a, 5a. Overexpression of the regulatory genes *PvMYB3A, PvMYB10B* and *P* separately or in combinations in pinto bean hairy roots revealed that the hairy roots overexpressing *P-PvMYB3A* fusion were able to significantly increase the expression of *PvDFR1A*, *PvDFR1B, PvANR2B* and *PvMATE8* compared to the control, which in turn increased the PA accumulation in those lines (Figure 5c, 5d).

The presence of *PvLAR1* transcripts in the vegetative tissues (leaf and stem) are reported for *M. truncatula* (Pang *et al*., 2007) and *L. japonicas* (Yoshida *et al*., 2008), however, these tissues do not accumulate catechin or PAs, which may indicate that there is not enough substrate present to produce catechin or that catechin is involved in another yet unknown biochemical reaction. It is possible that *PvLAR1*, like in *M. truncatula* and *L. japonicas*, is widely expressed in beans and synthesizes catechin or other metabolite depending on substrate availability. On the other hand, PvMYB3A root lines showed an increase in *PvLAR1* transcripts. A similar observation was made in strawberries (*Fragaria x ananassa*), where *LAR* expression was activated solely by the TT2-like MYB protein (Schaart *et al*., 2013), suggesting that at least one of the LBGs in the PA biosynthesis can be independently activated by the MYB. In common beans, epicatechin-3’-O-glucoside is transported into the vacuole by PvMATE8 (Islam *et al*., 2022). *PvMATE8* expression was elevated to a significantly greater level in P-PvMYB3A lines in pinto bean hairy roots, indicating that it has transcriptional regulation comparable to that of other LBGs.

ANR catalyzes the enzymatic reaction that leads to the generation of 2,3-*cis*-epicatechin as a PA starter unit from cyanidin in common bean (Coutin *et al*., 2017), Arabidopsis (Xu *et al*., 2015), *M. truncatula* (Li *et al*., 2016), peas (Hellens *et al*., 2010) and maize (Grotewold *et al*., 1994). ANR is also involved in the formation of 2,3-*cis*-leucocyanidin to serve as (−)-epicatechin extension units. This dual activity of ANR determines the compositional variation and level of PA accumulation across different plant species (Jun *et al*., 2021). Only PvMYB3A was able to bind to the *PvUANR2Bpro* among the three transcription factors P, PvMYB3A and PvMYB10B (Figure 6a). Notably, both P and PvMYB10B activated the *PvANR2B* gene expression (Figure 5c). Despite the presence of intact DNA-binding domains, P and PvMYB10B failed to bind the target promoter. In the trans-activation assay using *PvUANR2Bpro* driven *LUC* reporter gene expression, P exhibited a substantial increase in *LUC* expression only when combined with PvMYB3A (Figure 6c) demonstrating that P is not a DNA binding protein, but it influences transactivation through its interaction with PvMYB3A. This is a unique finding as no other bHLH tarnscription factor IIIf or TT8 subgroup with all functional domains exhibit non-DNA binding characteristics, while still functioning as a transcription activator (Heim *et al*., 2003). On the other hand, functional switch for the R (bHLH) in maize has been reported, where the DNA binding ability of the protein is determined by the presence of functional ACT-domains on the C-terminal end (Feller *et al*., 2006, Kong *et al*., 2012). Furthermore, plants have a small number of non-DNA binding bHLHs, the majority of which have been linked to light signaling pathways (Castelain *et al*., 2012). For example, Arabidopsis long hypocotyl in FR light (AtHFR1) a bHLH of VIIb subgroup, inhibits phytochrome-interacting factors 3, 4 and 5 (PIF3, PIF4 and PIF5) by forming non-DNA binding heterodimers in response to shade (Fairchild *et al*., 2000, Toledo-Ortiz *et al*., 2003, Hornitschek *et al*., 2009). Furthermore, the *PvUANR2Bpro* driven *LUC* gene expression was significantly reduced when P^sd^ rather than P was present in the MBW complex. This indicates the potential roles of the polymorphisms between *P* and *P^sd^*. (Islam *et al*., 2020).

Multiple MYBs are involved in the MBW complex to regulate PA biosynthesis across various plant species. For instance, in the seed coat epidermal tissues of Arabidopsis, MYB5 partially contributes to PA biosynthesis, even though TT2 is the primary R2R3-MYB (Gonzalez *et al*., 2009). Multiple MYBs with partially redundant roles have also been reported to be present in *M. truncatula* (Liu *et al*., 2014), *L. japonicus* (Yoshida *et al*., 2008), and other plants. However, in pinto bean, the presence of both PvMYB3A and PvMYB10B is not necessary for the transcription of *PvANR2B*, as evidenced by the trans-activation assays (Figure 6c). Furthermore, P doesn’t appear to have a preference for either of these PvMYBs. Since *PvMYB10B* (*J* gene), when non-functional leads to the ND phenotype (Erfatpour and Pauls, 2020), its strong correlation with seed coat darkening even in the absence of *PvUANR2B* promoter binding, indicates that it may be involved in controlling the early biosynthetic genes.

## Acknowledgements

We thank Alex Molnar, Kuflom Kuflu and Praveen Khatri (Agriculture and Agri-Food Canada, London, ON, Canada) for technical assistance. Special thanks to the undergraduate thesis students Madison McAvoy, Thebika Sivasri, Payton Barnes and Zilun Wang (Western University, London, ON, Canada) for their help. This research was supported by the Natural Sciences and Engineering Research Council of Canada Discovery Grant 04461-2018 RGPIN and Agriculture and Agri-Food Canada’s Abase J-001331 from to S.D.

## Supporting information

**Figure S1. Neighbour-joining trees of characterized DFR and ANR in legumes with common bean candidates.** Amino acid sequences of PvDFR and PvANR from *P. vulgaris* (landrace G199833) were aligned with characterised DFRs and ANRs from other plants. Phylogenetic trees were built in MEGAX with bootstrap values (%; 1000 replicates) shown next to the branch points. Putative PvDFR and PvANR are shown in grey. (a) DFR tree: Compared proteins from Arabidopsis AtDFR (AT5G42800), *G. max* GmDFR (ABM64803), *M. truncatula* MtDFR (AY389346), *Z. mays* ZmA1 (CAA28734), *Petunia x hybrida* PhDFR (MW929212), *L. japonicus* LjDFR (BAE19950) with PvDFRs. (b) ANR tree: Compared proteins Arabidopsis AtANR (AT1G61720), *G. max* GmANR (Glyma08g06630), and *M. truncatula* MtANR (AAN77735), *Fragaria x ananassa* FaANR (ABG76842), *Diospyros kaki* DkANR (BAF56654), *Gossypium hirsutum* GhANR (ABM64802), *Malus domestica* MdANR1 (AAZ17408), *Camellia sinensis* CsANR (AAT68773), *Theobroma cacao* TcANR (ADD51354), *Fagopyrum tataricum* FtANR (AHA14497), *Vitis vinifera* VvANR (DQ129684) with PvANRs.

**Figure S2. Stages of hairy root generation in common bean**. (1) callus formation at wounded sites at 5 days post-infection (dpi), (2) appearance of the first root from callus at 8-9 dpi, (3,4) roots growing at 11-14 dpi, (5-8) roots of length 1-2 cm ready to be transfered in a separate pot at 15 dpi by cutting 2 cm below the hairy roots, transplanted into a new pot, hairy roots were allowed grow over the vermiculite surface, (9) growth of hairy root at 18 dpi, (10) washed roots, (11) roots under a light microscope, (12) transgenic roots under UV-light showing GFP expression.

**Figure S3. Neighbour-joining trees of TT2 and TTG1 related MYB and WD40 proteins with common bean candidates.** Trees were built using MEGAX software. Bootstrap values (%; 1000 replicates) are shown next to the branch points. Protein sequence alignments were performed using ClustalW. Clustered candidates are shown in grey. (a) MYB tree: PvMYBs from *P. vulgaris* (landrace G199833) were compared with characterized TT2 like MYBs from other plants, Arabidopsis AtTT2 (AT5G35550.1), *M. truncatula* MtPAR (HQ337434), and *L. japonicus* LjTT2a (AB300033). (b) WD40 tree: PvWD40s from *P. vulgaris* (landrace G199833) were compared with TTG1 like WD40 proteins from other plants, Arabidopsis AtTTG1 (AT5G24520.1) *Z. mays* ZmPAC1 (AY115485), *P. hybrida* PhAN11 (U94748), *L. japonicas* LjTTG1 (AB490777), *M. truncatula* MtWD40-1 (EU040206).

**Figure S4. Expression analysis of PvMYB and PvWD40 candidates in different tissues of common bean.** Tissue-specific expression profile of the target genes in common bean. Transcript data retrieved from Phytozome for landrace G199833 was used for the generation of heatmap. The color key represents the relative transcript abundance in a row with the log_2_ transformed values. (a) MYB candidates, (b) WD40 candidates.

**Figure S5. Subcellular localization of the candidate interactors of P.** PvMYB3A, PvMYB10B, PvWD1A and PvWD9 were fused upstream of YFP reporter gene, co-transformed with NLS-CFP into *N. benthamiana*. Fluorescence were visualized in leaf epithelial cells by confocal microscopy. Merged signal was collected by sequential scanning of YFP and CFP channels. Scale bar=30 μm.

**Figure S6. Protein-protein interaction between candidate proteins involved in the MBW complex formation in common bean.** BiFC assay between proteins was assayed by co-expression of translational fusions with N–terminal (YN) and C–terminal (YC) fragments of YFP in *N. benthamiana*. Proximity of the two fragments results in a functioning fluorophore. Scale bar= 40 µM.

**Figure S7. Functional analysis of PvMYB3A, PvMYB10B and PvWD9 using Arabidopsis mutants.** (a) DMACA stained seed of WS2, *tt2*, *tt2* transformed lines with PvMYB10B, *tt2* transformed lines with PvMYB3A, (b) Genetic complementation of *ttg1* in Arabidopsis by variants of PvWD9. Phenotypes of Columbia-0, *ttg1* and *ttg1* transformed lines with PvWD9 are shown. From top, photographs of DMACA stained seeds, Anthocyanin in seedlings, trichrome in young leaf, Ruthenium red stained seeds showing seed mucilage and root hair patters.

**Figure S8:** Sequence alignment of *PvANR2B* (cultivar G19833) and *PvUANR2B* (pinto cultivar UI111) promoter region (−2000 bp).

**Table S1.** Oligonucleotides used in this study.

**Table S2.** *Cis*-acting elements in *PvUANR2B* promoter

## Data Availability Statement

The data that support the findings of this study are available on request from the corresponding author.

